# Phase transitions may explain why SARS-CoV-2 spreads so fast and why new variants are spreading faster

**DOI:** 10.1101/2021.02.16.431437

**Authors:** J. C. Phillips, Marcelo Moret, Gilney F. Zebende, Carson C. Chow

## Abstract

The novel coronavirus SARS CoV-2 responsible for the COVID-19 pandemic and SARS CoV-1 responsible for the SARS epidemic of 2002-2003 share an ancestor yet evolved to have much different transmissibility and global impact^1^. A previously developed thermodynamic model of protein conformations predicted that SARS CoV-2 is very close to a thermodynamic critical point, which makes it highly infectious but also easily displaced by a spike-based vaccine because there is a tradeoff between transmissibility and robustness^2^. The model identified a small cluster of four key mutations of SARS CoV-2 that promotes much stronger viral attachment and viral spreading. Here we apply the model to two new strains (B.1.1.7 and B.1.351)^3^ and predict, using no free parameters, how the new mutations can further enhance infectiousness.

## Introduction

The SARS-CoV-2 spike (S) is a glycosylated trimeric class 1 fusion protein that enables the virus to enter cells^4,5^. The S glycoprotein rests in a metastable prefusion state that must undergo conformational changes before the virus can fuse to the cell membrane. Given the importance of dynamics for S function it can be better understood as a thermodynamic object immersed in water. This hypothesis has been validated by a confirmed prediction that vaccines based on S would be very effective and finding four key mutations that enhance viral attachment and infectiousness^2^. This is nontrivial as vaccine efficacy is not always guaranteed^6^.

It has long been hypothesized that some biological systems including proteins may extract important functional benefits from operating at the edge of instability, halfway between order and disorder, i.e., in the vicinity of the critical point of a phase transition^7–9^. Topological constraints, namely, the importance of protein inside and outside shape extremals as measured by the interactions of amino acid side groups with water are paramount^8,10^. Each S chain consists of over 1200 amino acid residues, with approximately 300 of these having mutated from SARS-CoV-1 (CoV-1) to SARS-CoV-2 (CoV-2)^11^. Here, we show how the nine additional S amino acid mutations in the B.1.1.7 variant can lead to a small increase in spreading rate while not diminishing the efficacy of existing S-based vaccines. However, the additional mutations in B.1.351 may diminish vaccine efficacy somewhat.

## Materials and Methods

The model utilizes a thermodynamic amino acid scale^12^ that considers the loss of solvent-accessible surface area (ASA) surrounding a central residue. Moret and Zebende^12^ found in over 5000 different protein segments that the ASA for a specified amino acid residue at the center of a protein fragment scales as a power-law of the length of the fragment with a well-ordered negative exponent for all 20 amino acids (See Fig. 1 for a schematic of the analysis and Table 1 for the values). The existence of such universal amino acid–water interaction parameters is by itself convincing evidence for the thermodynamic phase transition model^7,9^. The average accuracy of each of the 20 exponents is better than 1% ^2,12^. In the absence of a phase transition critical point, the likelihood that 20 such accurate power-law exponents would coexist independently is astronomically small (less than 10^−40^). To our knowledge, proteins are the only large-scale networks that exhibit both first-order unfolding phase transitions and second-order conformational phase transitions described by fractals^2,12^.

**Table 1.**
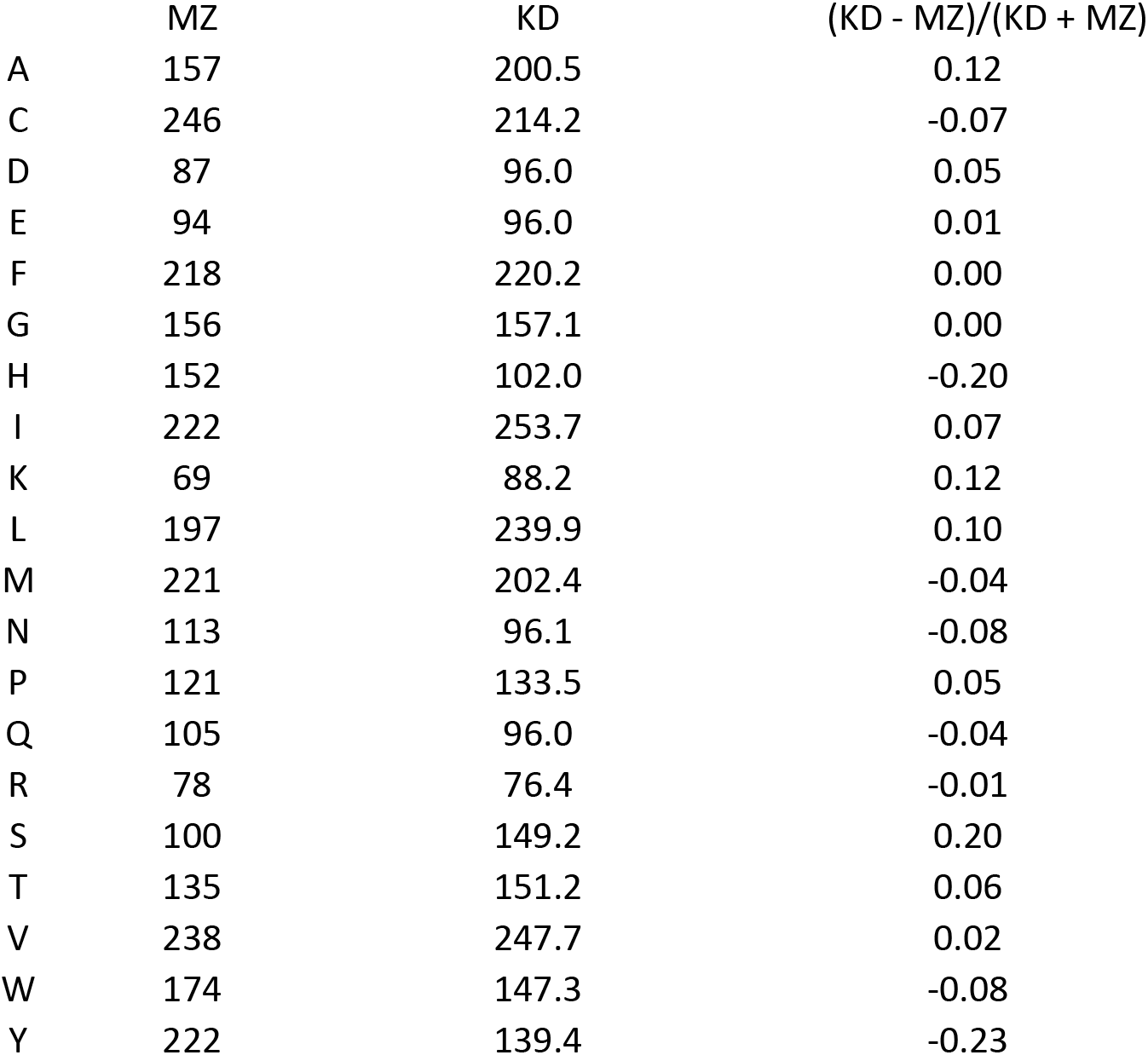
The shifted and rescaled hydropathic values Ψ for the Moret Zebende (MZ) and Kyte-Doolittle (KD) scales. The overall correlation is 86%, and in practice the differences are large enough to be reflected in protein function. For different proteins, one or the other of the two scales is better. Here and previously^2^ only the MZ scale gives synchronous edges that optimize conformational changes for faster spreading for CoV-2 and its mutants.

**Table 2.** Scores for main hydrophilic edges (based on CoV-2 sites). Taking the four most hydrophilic edges (1,2,4,6), the coefficient of variation for the hydropathy scores are 0.04, 0.01, 0.007, and 0.01 for CoV-1, CoV-2, B.1.1.7, and B.1.351, respectively. There was a large decrease between CoV-1 and CoV-2 and a smaller one between CoV-2 and B.1.17, indicating a progression towards a symmetric critical point. Note that new mutations can be significant if they lie within 17 amino acids of these five and involve large (non-synonomous) changes in Ψ(aa). An example of a nonsynonomous change is A to D (change 70), while the E to K change is only 8. (see Suppl. Table 1)

| Strain \ Edges | 1 (454) | 2 (569) | 3 (694) | 4 (803) | 5 (952) | 6 (1156) |
| --- | --- | --- | --- | --- | --- | --- |
| CoV-1 | 137.9 | 131.3 | 146.5 | 142.5 | 137.5 | 137.5 |
| CoV-2 | 137.1 | 137.7 | 140.1 | 135.2 | 138.3 | 135.2 |
| B.1.1.7 | 137.1 | 135.7 | 141.0 | 135.2 | 138.3 | 135.2 |
| B.1.351 | 137.1 | 137.7 | 141.7 | 135.2 | 138.3 | 135.2 |

**Table 3.** Scores for a selection of hydrophobic edges (based on CoV-2 sites). While most of the hydrophobic peaks are similar between CoV-2 and the new variants, there was a significant increase in hydrophobicity for peak B in B.1.351.

| Strain<br>Edges | A<br>(131) | B<br>(227) | C<br>(378) | D<br>(496) | E<br>(607) | F<br>(755) | G<br>(894) | H<br>(1116) | I<br>(1231) |
| --- | --- | --- | --- | --- | --- | --- | --- | --- | --- |
| CoV-1 | 164.7 | 164.1 | 166.2 | 171.0 | 159.3 | 152.1 | 168.4 | 163.4 | 192.0 |
| CoV-2 | 167.6 | 159.9 | 164.2 | 169.8 | 166.7 | 167.2 | 168.3 | 165.3 | 196.2 |
| B.1.1.7 | 165.2 | 159.9 | 164.2 | 172.9 | 166.7 | 167.2 | 168.3 | 167.1 | 196.2 |
| B.1.351 | 167.6 | 163.1 | 164.2 | 172.7 | 168.6 | 167.2 | 168.3 | 165.3 | 196.2 |

**Fig. 1.**
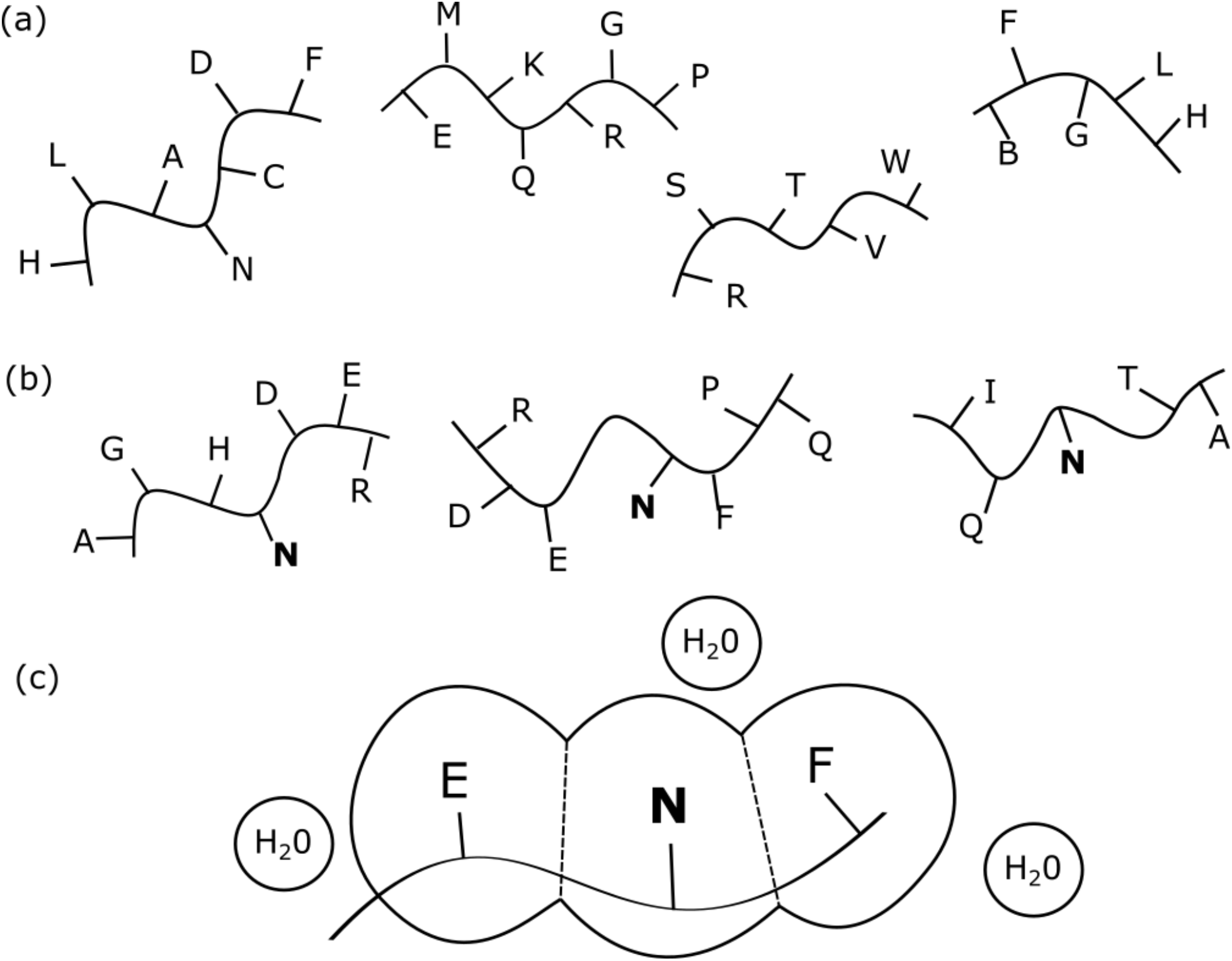
(a) Moret and Zebende^12^ began with a set of 5526 protein segments of varying length M up to 45. A few short examples are shown here, with their amino acid side chains. (b) These segments were grouped into 20 subsets. Each subset has the same amino acid at its center, so each subset contains about 250 segments. (c) The amino acids side chains are each surrounded by a sphere with radius set by van der Waals interactions. Where the spheres overlap, they are cutoff by planes equidistant from their chain contacts (dotted lines). The surface area of the central amino acid that is accessible to water molecules is then calculated. These surface areas are then plotted for each amino acid as functions of chain length M. Increasing M decreases surface areas as the chains fold back upon themselves. These decreases were fitted well by power-laws; such power-laws are known to be characteristic of fractal structures and second-order phase transitions very close to critical points^7,8^.

The 20 exponents give a measure of the average hydropathy of each residue at the center of an arbitrary background neighborhood. The smaller the magnitude of the exponent, the more hydrophilic the amino acid will be on average. This differs from the many attempts to assign residue the more likely it will reside near the outside of the protein and *vice versa* for the more hydropathy based on chemical properties of an amino acid in isolation^12^. The more hydrophilic a hydrophobic. However, given that the residues are situated sequentially on a chain, the effect of neighboring residues must be taken into account. This requires deducing an effective domain size that dominates the protein’s conformation, whose details at the molecular level are unknown. This seemingly intractable task is circumvented with the conjecture that natural selection favors sequences such that the protein is close to a thermodynamic critical point. Evidence for positive selection has been demonstrated in SARS-CoV-2^13^. Nearer the critical point, it can function more reversibly and at lower temperatures^7^. In this case, there could be one dominant domain length and this conjecture has been applied successfully to many proteins^10,14^. The model also posits a mechanism for the increase in infectiousness of CoV-2 based on synchronized attachment^2^. Of the over 300 mutations between CoV-1 and CoV-2, the model predicts that only four key mutations affected its infectiousness. The new B.1.1.7 variant, which contains only 9 amino acid mutations (3 deletions and 6 substitutions)^3^, seems to be transmitting at a higher rate than CoV-2^15^. We show how a single additional key mutation has increased the transmissibility of B.1.1.7, and what this means for future increases and vaccine efficacy.

The predictions are made by computing the coarse-grained (shifted and rescaled, see Table S1) magnitude of the exponent along S over a domain length W to create a matrix of hydropathy scores Ψ(R,W), where a larger value means that the residue at site R acts more hydrophobically. W is the width of a sliding window centered on each R to average nearby interactions. Ψ(R,W) oscillates between hydrophilic to hydrophobic values with decreasing amplitude as W increases. The variance of Ψ(R,W) is thus a roughness measure of the protein, and an optimal W can be selected that minimizes the ratio of variances between proteins in a family, which is often connected to observed static structural features^8^. The successful analysis of CoV-1/CoV-2 differences^2^ used W=35, so that choice is used here as well, to analyze the quite small B.1.1.7 additional mutational differences, using no new or free parameters. In particular, the conclusion reached before^2^ that the spike attachment dynamics are synchronized by hydrophilic extrema is preserved.

## Results

Figure 2 shows the profile of Ψ(R,35) for CoV-2 S together with the variants B.1.1.7 and B.1.351. The notable features include a large hydrophobic maximum near residue 1231 and 6 deep minima between 400 and 1200. The large maximum is located in the virus transmembrane domain. The minima represent edges of hydrophilic regions. Edge 1 (454) is located in the Receptor Binding Domain (RBD), 2 (569) in C-Terminal Domain 1 (CTD1), 3 in the Linker between S1 and S2 (694), 4 in the Fusion peptide in S2 (803), 5 (952) in heptad repeat 1 (HR1), and 6 (1156) in the Linker to the stem of S2^16^. Although these edges do not necessarily require to be associated with any observable structural features, they seem to be located in salient regions for dynamics. Edges 1, 2, 4, and 6 are the most hydrophilic. They have similar magnitudes and are much more similar than the corresponding edges of CoV-1 (see Table 1). This symmetry is a signature of a critical point and lends the hydrophilic domains more conformational mobility, particularly for acting in synchrony. The symmetry is also fragile and can be easily broken by mutations as confirmed by mutation simulations (data not shown). The differences between Ψ(R,35) for CoV-2 and B.1.1.7 are small, but close inspection of the hydrophilic edges shows that edges 1, 2, 4 and 6, which were nearly equal already in CoV-2, have become even more equal in B.1.17 (see Table 1). The coefficient of variation (C.V.) has decreased from 0.01 to 0.007. This decrease occurs because edge 2 has shifted into better agreement with edges 4 and 6. Such long-range or allosteric interactions occur in motor proteins, where they were quantified by hydropathic scaling^17^. They are known to occur in principle^18^ but when small but are detectable using the hydropathic scale with 20 exponents. Equalization of hydrophilic edges was recognized as the cause of domain attachment synchronization in the evolution of CoV-1 to CoV-2 for second, but not first, order phase transitions^2^. The observed differences between upper respiratory tract (e.g. throat) and lower respiratory tract (e.g. lung) infection are consistent with our model^19^.

**Fig. 2.**
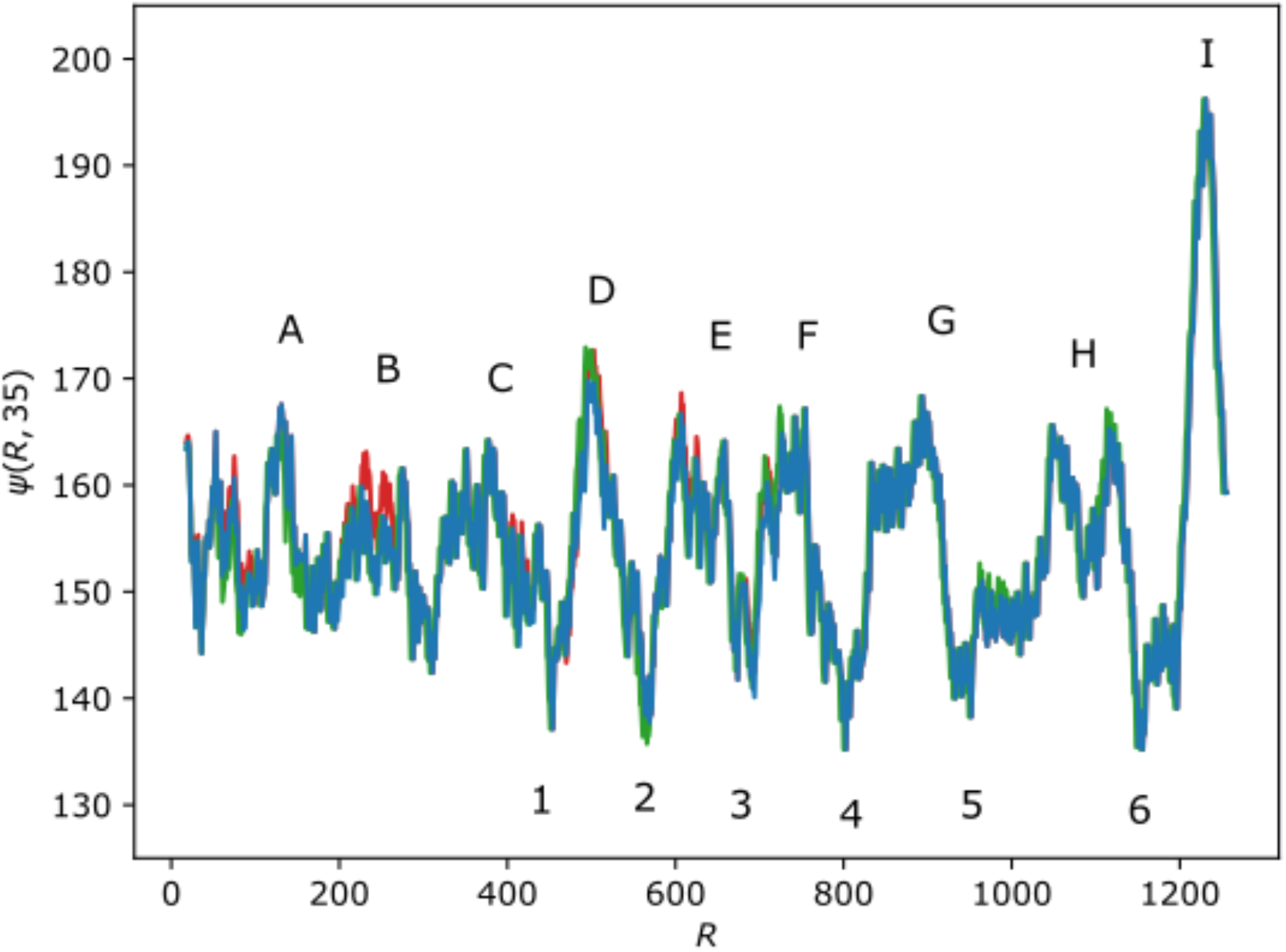
Hydropathy score Ψ(R,W=35) for SARS-CoV-2 S (blue), B.1.1.7 (green), and B.1.351 (red). Hydrophilic edges 1 through 6 are labeled. Edges 1 and 2 are nearly symmetric as are edges 4 and 6 (see Table 1). In SARS-CoV-1, none of the edges in close proximity were symmetric but in B.1.1.7, edge 2 has evolved to become more symmetric with edges 4 and 6, while 5 remains the same. Hydrophobic edges are labeled A through I. Edge B is more hydrophobic and more symmetric with edge C in B.1.351 than SARS-CoV-2.

The South Africa strain B.1.351^19^ has eight mutations, L18F, D80A, D215G, R246I, K417N, N501Y, and A701V. It also includes D614G, which predates the strain. These mutations do not affect the hydrophilic edges compared to CoV-2, but they do make the hydrophobic peak located at residue 227 larger, which is in the N-terminal domain adjacent to the the receptor binding domain. This increased hydrophobicity could stabilize the virus in aerosols^20^. The near leveling of the hydrophobic peak near 227 with the receptor binding peak near 380 suggests that the two peaks could bind together to aerosol surfaces. Overall, there seems to be a qualitative difference in the mechanisms used by the B.1.17 and B.1.351 to supplant CoV-2. This could be reflected in the dependence of their relative selectivity on prevailing control measures and could stabilize this strain against some vaccines.

## Discussion

The higher transmissibility of B.1.1.7 compared to CoV19 is mostly caused by the single key mutation A570D. At first this appears to be a cause for concern. Could more key mutations bring the edges into better agreement, increasing transmission further? This has low probability as it would require coordinated hydrophilic mutations within the narrow range of 35 residues surrounding an edge. Additionally, the great success of vaccines based on S was predicted because even a small disturbance in S will tend to drive it away from the critical point^2^. For the virus to attach to a cell, S must act in a coordinated fashion and this is impaired away from the critical point. Thus, the S-based vaccines that are already available are expected to be equally successful not only for B.1.1.7, but for any future mutation as well that increases transmissibility by moving the virus still closer to its critical point. B.1.351 may be deploying a different strategy by leveling two hydrophobic peaks to stabilize S, which may explain why vaccine efficacy differs between the two new strains.

All S mutations are easily evaluated using our measure, which can also be used to design animal experiments to test S mutations for their nearness to criticality and thus transmissibility. Our score evaluates transmissibility due to efficacy of viral attachment to cells but does not address the effect of mutations that may alter mechanisms after the viral material has entered the cell.

Finally the predictions (with no new parameters) were made possible through experience gained from the many previous studies that utilized the global 21^st^ century protein database^21^.

## References

1. Lu, R. et al. Genomic characterisation and epidemiology of 2019 novel coronavirus: implications for virus origins and receptor binding. The Lancet 395, 565–574 (2020).

2. Phillips, J. C. Synchronized Attachment and the Darwinian Evolution of Coronaviruses CoV-1 and CoV-2. ArXiv200812168 Q-Bio (2020).

3. Preliminary genomic characterisation of an emergent SARS-CoV-2 lineage in the UK defined by a novel set of spike mutations. Virological https://virological.org/t/preliminary-genomic-characterisation-of-an-emergent-sars-cov-2-lineage-in-the-uk-defined-by-a-novel-set-of-spike-mutations/563 (2020).

4. Wrapp, D. et al. Cryo-EM structure of the 2019-nCoV spike in the prefusion conformation. Science 367, 1260–1263 (2020).

5. Walls, A. C. et al. Structure, Function, and Antigenicity of the SARS-CoV-2 Spike Glycoprotein. Cell 181, 281–292.e6 (2020).

6. Mazur, N. I. et al. The respiratory syncytial virus vaccine landscape: lessons from the graveyard and promising candidates. Lancet Infect. Dis. 18, e295–e311 (2018).

7. Muñoz, M. A. Colloquium: Criticality and dynamical scaling in living systems. Rev. Mod. Phys. 90, 031001 (2018).

8. Phillips, J. C. Fractals and self-organized criticality in proteins. Phys. Stat. Mech. Its Appl. 415, 440–448 (2014).

9. Phillips, J. C. Thermodynamic Scaling of Interfering Hemoglobin Strain Field Waves. J. Phys. Chem. B 122, 9324–9330 (2018).

10. Phillips, J. C. Scaling and self-organized criticality in proteins: Lysozyme c. Phys. Rev. E 8.

11. Severe acute respiratory syndrome coronavirus 2 (2019-nCoV) (SARS-CoV-2). https://www.uniprot.org/taxonomy/2697049.

12. Moret, M. A. & Zebende, G. F. Amino acid hydrophobicity and accessible surface area. Phys. Rev. E 75, 011920 (2007).

13. Berrio, A., Gartner, V. & Wray, G. A. Positive selection within the genomes of SARS-CoV-2 and other Coronaviruses independent of impact on protein function. PeerJ 8, (2020).

14. Phillips, J. C. Darwinian Evolution of Taste. ArXiv200812984 Q-Bio (2020).

15. Davies, N. G. et al. Estimated transmissibility and severity of novel SARS-CoV-2 Variant of Concern 202012/01 in England. 35.

16. Cai, Y. et al. Distinct conformational states of SARS-CoV-2 spike protein. Science 369, 1586–1592 (2020).

17. Phillips, J. C. Self-organized networks: Darwinian evolution of dynein rings, stalks, and stalk heads. Proc. Natl. Acad. Sci. 117, 7799–7802 (2020).

18. Marvin, J. S. et al. The rational design of allosteric interactions in a monomeric protein and its applications to the construction of biosensors. Proc. Natl. Acad. Sci. 94, 4366–4371 (1997).

19. Tegally, H. et al. Emergence and rapid spread of a new severe acute respiratory syndrome-related coronavirus 2 (SARS-CoV-2) lineage with multiple spike mutations in South Africa. medRxiv 2020.12.21.20248640 (2020) doi:10.1101/2020.12.21.20248640.

20. Jayaweera, M., Perera, H., Gunawardana, B. & Manatunge, J. Transmission of COVID-19 virus by droplets and aerosols: A critical review on the unresolved dichotomy. Environ. Res. 188, 109819 (2020).

21. Guttmacher, A. E. & Collins, F. S. Welcome to the Genomic Era. N. Engl. J. Med. 349, 996–998 (2003).

